# Novel Lipophagy Inducers as Potential Therapeutics for Lipid Metabolism Disorders

**DOI:** 10.1101/2025.03.11.642628

**Authors:** Rachel Njeim, Bassel Awada, Haley Donow, Haley Gye, Cole Foster, Colin Kelly, Judith Molina, Sandra Merscher, Marcello Giulianotti, Alessia Fornoni, Hassan Al-Ali

## Abstract

Dysregulation of lipid homeostasis is associated with a wide range of pathologies encompassing neurological, metabolic, cardiovascular, oncological, and renal disorders. We previously showed that lipid droplet (LD) accumulation in podocytes contributes to the progression of diabetic kidney disease (DKD) and reducing LDs preserves podocyte function and prevents albuminuria. Here, we sought to identify compounds that treat pathological LD accumulation. We developed a phenotypic assay using human podocytes and deployed it to screen a combinatorial library comprising over 45 million unique small molecules. This led to the identification of a compound series that effectively reduce LD accumulation in stressed podocytes. Mechanistic studies revealed that these compounds activate lipophagy, reduce LD accumulation, and rescue podocytes from cell death. In contrast, compounds known to induce general autophagy failed to mimic these effects, indicating a novel lipophagy-specific mechanism of action (MoA), which was confirmed with unbiased phenotypic profiling. An advantage of this therapeutic strategy is its potential to not only halt the progression of pathological lipid accumulation but also reverse it. These compounds will serve as tools for uncovering novel drug targets and therapeutic MoAs for treating DKD and other diseases with similar etiologies.

**GRAPHICAL ABSTRACT:** 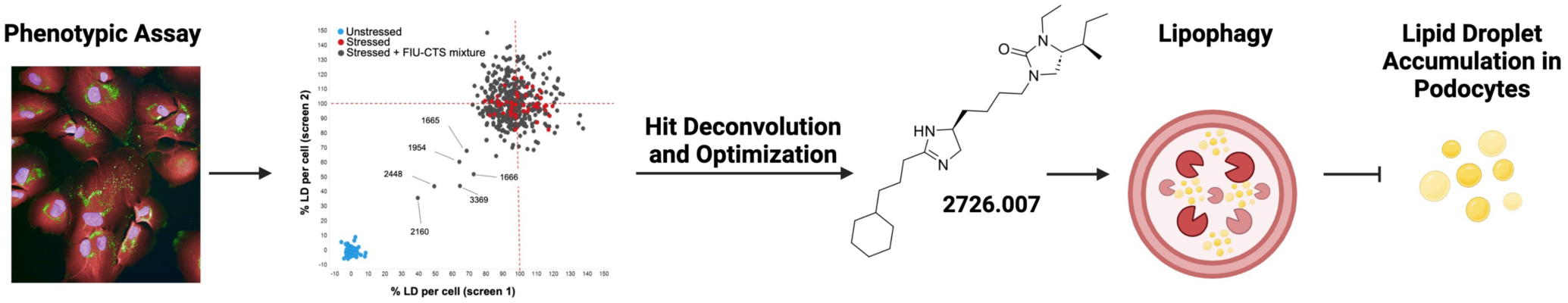

## INTRODUCTION

Lipid droplets (LDs), once thought to be mere subcellular storage units, are now recognized as crucial cellular organelles with key physiological functions^1^. These droplets consist of a hydrophobic core made primarily of neutral lipids, including triglycerides (TG) and cholesterol esters (CE), surrounded by an amphipathic phospholipid monolayer^2^. LD accumulation has been shown to initially mitigate lipotoxicity caused by endoplasmic reticulum stress and reactive oxygen species, while also supporting the sequestration, trafficking, and eventual export of harmful cytotoxic components. However, disruptions in the processes can lead to counterproductive LD accumulation and contribute to the development of various diseases including neurological, metabolic, cardiovascular, oncological, and renal disorders^1^.

Extensive research points to a central role of LD accumulation in the development and progression of Diabetic Kidney Disease (DKD)^3,4^, a leading cause of end-stage renal disease (ESRD). DKD affects approximately 40% of individuals with diabetes^5^. In the U.S., an estimated 35.5 million people have Chronic Kidney Disease (CKD), with diabetes and hypertension remaining the two most common causes^6^. DKD is characterized by persistent albuminuria and a gradual decline in glomerular filtration rate, with histopathological features including podocyte injury, tubular and glomerular basement membrane thickening, mesangial expansion, and glomerulosclerosis^7^. Kidney biopsies from patients with DKD demonstrate downregulation of genes involved in fatty acid β-oxidation, including peroxisome proliferator-activated receptor alpha (PPAR-α), carnitine palmitoyltransferase 1, acyl-CoA oxidase, and liver fatty acid-binding protein (L-FABP), as well as genes involved cholesterol efflux, including ATP-binding cassette A1 (ABCA1), ATP-binding cassette G1 (ABCG1), and apolipoprotein E (ApoE). Conversely, receptors involved in cholesterol uptake, such as the low-density lipoprotein receptor (LDLR), oxidized LDLR, and acetylated LDLR, are upregulated^4^.

Consistent with these findings, we observed increased LD accumulation in cultured podocytes exposed to sera from patients with DKD and in glomeruli from individuals in the early stages of diabetes^4^. LD accumulation in podocytes is associated with increased cardiolipin peroxidation, mitochondrial dysfunction, and podocyte injury^8^, ultimately predisposing podocytes to cell death. Notably, while sequestering cholesterol with cyclodextrin (CD) protects podocytes from actin cytoskeleton remodeling and cell blebbing, blocking cholesterol synthesis with statins did not prevent these changes^9^. Furthermore, we demonstrated that podocyte-specific ABCA1 deficiency exacerbated TNF-induced albuminuria, whereas ABCA1 overexpression or CD treatment reduced albuminuria and slowed DKD progression in a mouse model with podocyte-specific NFATc1 activation, mimicking DKD-associated glomerulosclerosis^10^. Our previous research also showed that podocyte LD accumulation contributes to the progression of non-metabolic kidney diseases, such as focal segmental glomerulosclerosis (FSGS) and Alport Syndrome^11^. Treatment with small molecule compounds that promote ABCA1-dependent cholesterol efflux^12^ or 2-hydroxy-b-cyclodextrin reduced renal CE, LDs, and cholesterol crystal content in experimental AS and FSGS^11^. Importantly, LD accumulation in podocytes occurred independently of serum lipid levels^10,13^, emphasizing that this process is cytopathological rather than a passive consequence of elevated systemic lipid levels^14^. Consequently, drugs that lower systemic lipids through the inhibition of 3-hydroxy-3-methylglutaryl coenzyme A (HMG-CoA) reductase do not reduce LD accumulation in kidney cells^15^ and fail to prevent DKD progression in patients^16,17^. Therefore, developing molecules that directly target LD accumulation may be essential for treating DKD and other disorders associated with this pathological process.

In this study, we developed a robust phenotypic assay to assess LD accumulation in human podocytes^18^. We screened a mixture-based library comprising over 45 million unique small molecules and identified a novel class of PAINS-free compounds that effectively protect podocytes from LD accumulation, lipotoxicity, and cell death. Structure-activity relationship (SAR) studies led to the development of 2726.007, an analog with improved properties. Mechanistic studies revealed that 2726.007 protects podocytes by inducing lipophagy and reducing inflammatory signaling, potentially offering broader clinical impact in the context of pathologies mediated by LD accumulation, which are often characterized by chronic inflammation^11,19,20^.

## RESULTS AND DISCUSSION

### Phenotypic assay for quantifying LD accumulation in stressed human podocytes

We previously developed a robust phenotypic assay for quantifying LDs in differentiated human podocytes^18,21^. Phenotypic screening offers a significant advantage in drug discovery, enabling the identification of biologically active compounds without prior knowledge of specific targets or mechanisms of action. This is particularly relevant in nephrology, where assays that faithfully capture pathobiology are lacking and target-based approaches have been less efficient^22^.

We sought to develop a clinically relevant phenotypic screening approach with differentiated human podocytes to increase the translatability of discovered hits. Given that sera from well-characterized DKD patients are not available in sufficient quantities for primary screening and can only be used in secondary screening and follow-up studies, we sought to identify an alternative stressor for our primary assay that meets three key criteria: 1) induce relevant and sufficient LD accumulation to give the assay a suitable Z-factor, 2) be readily scalable for screening purposes, and 3) be appropriate for discovering compounds that modify disease induced by clinical samples (e.g. DKD patient sera). To address this, we tested various podocyte stressors known to be relevant to DKD, including tumor necrosis factor (TNF), free fatty acids of various chain lengths, oxidized and unoxidized LDL, and free cholesterol. We also tested stressors relevant to other kidney disorders, such as collagen I, which is implicated in podocyte lipotoxic injury in Alport Syndrome^23^. While most of these factors induced LD accumulation, none yielded a Z-factor > 0.5. We then proceeded to test chemical inducers of podocyte stress, with puromycin, a known inducer of podocyte stress^24^, emerging as the most effective for inducing LD accumulation in human podocytes.

Under unstressed conditions, podocytes exhibit minimal LD accumulation **(Fig. 1A, 1B)**. However, treatment with puromycin (30 µg/mL) leads to significant LD accumulation **(Fig. 1A, 1B)** accompanied by a decrease in cellular viability **(Fig. 1C)**. Notably, the Z-factor for this assay consistently exceeded 0.5 **(Fig. 1B)**, highlighting its robustness and suitability for screening^25^.

**Figure 1:**
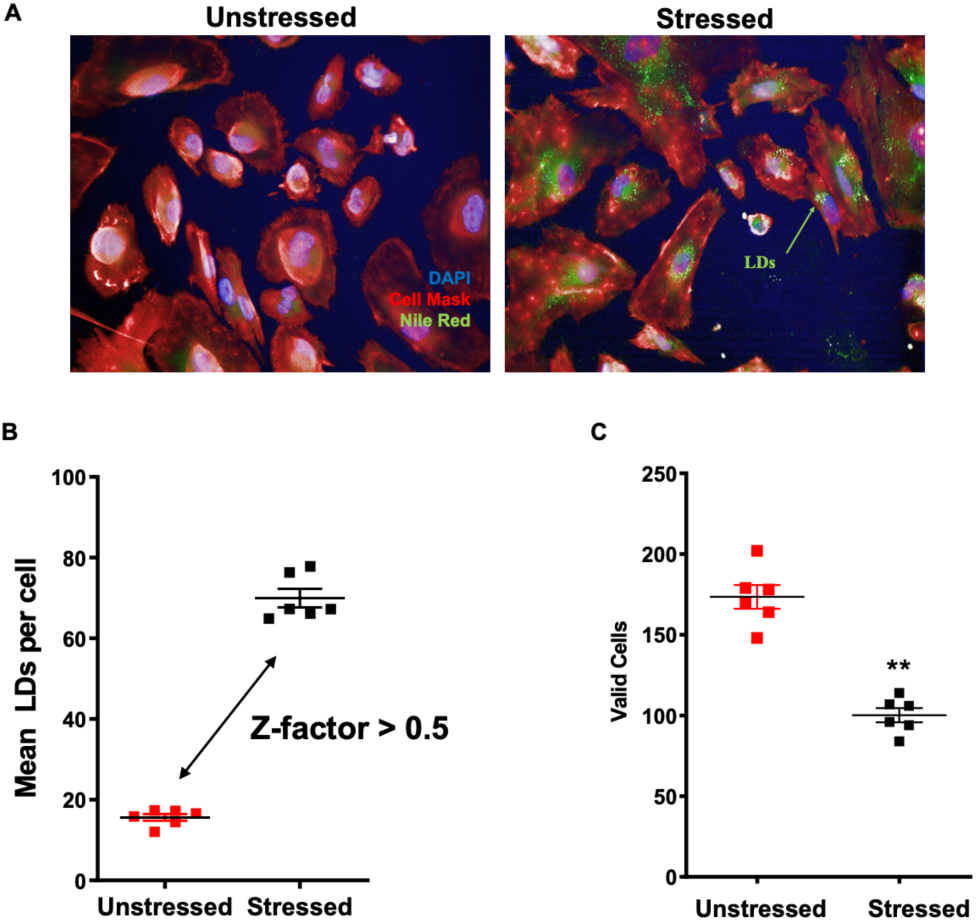
Phenotypic assay for LD quantification in human podocytes using the Opera Phenix HCS system. (A) Representative single-field OPERA confocal images of differentiated podocytes stained with DAPI (nuclei, blue), Nile Red (LDs, green), and CellMask Deep Red (cell bodies, red). Podocytes were treated for 48 hours and analyzed for (B) mean LD per cell and (C) valid cell count. Data represent mean ± SEM.

### Hit identification by scaffold ranking and positional scanning

We deployed the phenotypic assay to screen the Florida International University Center for Translational Science (FIU-CTS) scaffold ranking library, which consists of compounds that are systematically arranged in positional scanning format^26–30^. The diversity of the FIU-CTS lead generation library has been characterized and described quantitatively using molecular scaffolds, molecular properties, and structural fingerprints^31,32^. The library provides unique lead compounds suitable for optimization and advancement to preclinical and clinical studies, and its successful use has been reviewed^33,34^. The superior performance of the FIU-CTS library in comparison to the Prestwick Chemical Library (PCL), MLSMR, and a computational screen was previously reported^35^. Mixtures were tested at four different concentrations (5, 1, 0.2, and 0.04 µg/ml) to identify molecular scaffolds that reduce LD accumulation in stressed podocytes. The primary screen revealed several potential hits, including 1665, 1666, 1954, 2160, 2448, and 3369 **(Fig. 2A)**. Mixtures 1665 and 2448 are both polyamines and synthesized separately using slightly different reagents. Their similar activity in this screen highlights the robustness of both the chemistry and the assay. Mixtures 1666, 1954 and 2160 are derived from piperazine, while mixture 3369 is derived from dihydroimidazolyl-butyl-cyclic thiourea **(Fig. 3)**. We confirmed mixtures 2448, 2160, 1954, and 3369 by retesting with fresh stocks **(Fig. 2B)**. In our experience, certain compounds can alter LD numbers without affecting overall lipid content, potentially by modulating the enzymes responsible for merging LDs into larger droplets, which exhibit stronger fluorescence^36^. Therefore, hits that increase LD intensity were disqualified **(Fig. 2C)**. Additionally, since this is a loss-of-signal assay, we filtered out nuisance, typically cytotoxic compounds^37,38^. Valid cell counts (i.e. number of viable cells at endpoint) were tracked and normalized to the counts in the unstressed condition **(Fig. 2D)**. These two filters disqualified mixtures 2448, 2160, and 1954, thereby leaving only 3369 as a true hit. Importantly, 3369’s ability to prevent LD accumulation was associated with improved podocyte health in a dose-dependent manner **(Fig. 2D)**. To verify the disease-modifying activity of 3369, we tested it in a secondary assay using sera from patients with type 1 diabetes and kidney disease from the Finnish Diabetic Nephropathy (FinnDiane) Study as an inducer of podocyte stress and injury^9^. Our findings demonstrate that 3369 markedly prevented podocyte LD accumulation induced by DKD sera **(Fig. 2E)**.

**Figure 2:**
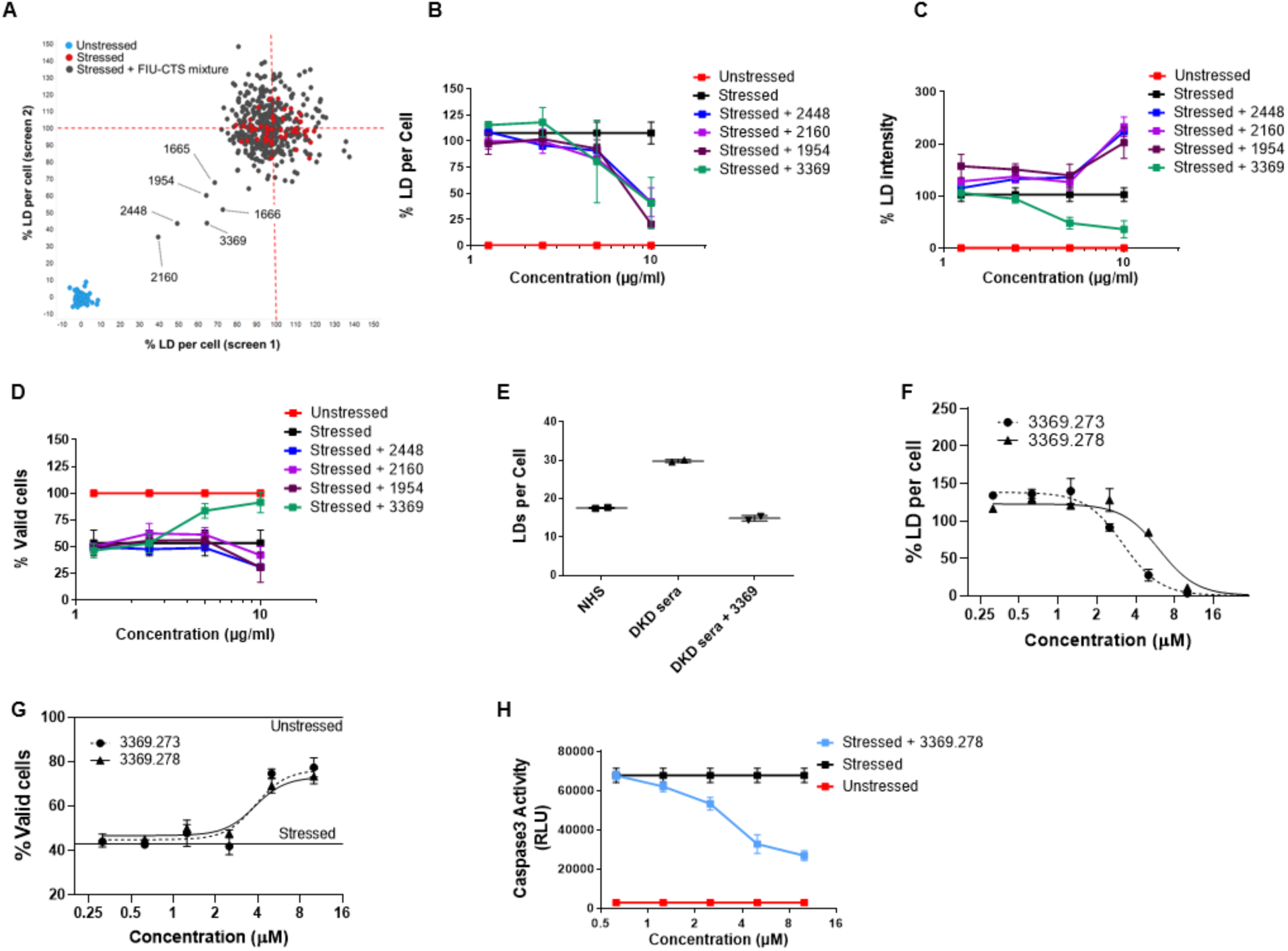
Hit discovery and confirmation in the primary screen. Scaffold ranking screen of Florida International University Center for Translational Science (FIU-CTS) collection: A) High content analysis of stressed podocytes screened with the FIU-CTS mixture library at four different concentrations (5, 1, 0.2, and 0.04 µg/ml). The scatter plot depicts results from two separate runs, whereby data are normalized to the means of stressed (set to 100%) and unstressed (set to 0%) conditions in each screen. Validation of active mixtures: (B) mean LD per cell (averaged per well) normalized to stressed condition (% LD per cell), (C) mean LD intensity normalized to stressed condition (% LD Intensity), and (D) valid cell count normalized to unstressed condition (% valid cells). Data represent mean ± SD, n=4 technical replicates. (E) Podocytes were treated with normal human sera (NHS) (4 % v/v) or DKD patient sera co-treated with either vehicle or 3369 (1 µg/ml) and assayed for LD accumulation. Data represent mean ± SD, n=2 technical replicates. Positional scanning and validation of individual hits from the 3369 mixture: (F), (G) Dose-response testing of 3369.278 and 3369.273 in the primary assay. Data represent mean ± SD, n=3 technical replicates. (H) Apoptosis assessed by caspase 3 activity. Data represent mean ± SD, n=3 technical replicates.

**Figure 3:**
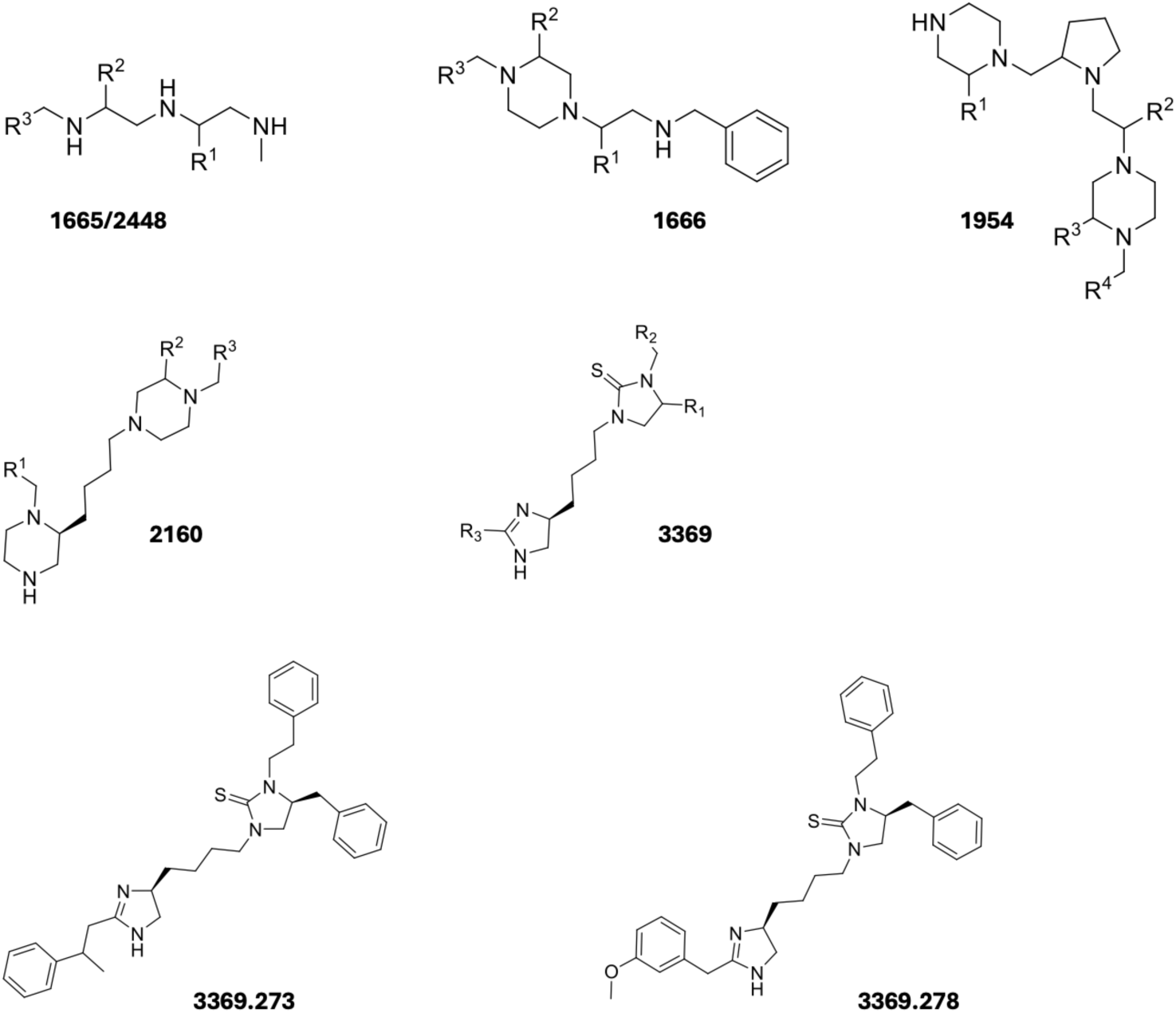
Chemical structures of initial hits. Shown are the structures for mixtures 1665, 2448, 1666, 1954, 2160, and 3369 as well as the two initial hit individual compounds 3369.273 and 3369.278. For the mixture samples the “R” groups represent positions where a diversity of functionalities has been incorporated into the sample. Mixtures 1665 and 2448 are two different synthesis lots of the sample mixture, screening both samples allows us to test the robustness and repeatability of the chemistry and assays.

Next, we proceeded to perform hit deconvolution from this mixture as previously described^34^. We synthesized and HPLC-purified predicted hits and assessed their activity in the phenotypic assay. All analogs exhibited varying levels of activity, revealing a rich SAR landscape and high feasibility for hit-to-lead optimization. Representative compounds, 3369.278 and 3369.273 **(Fig. 3)**, demonstrated potent inhibition of LD accumulation while restoring podocyte numbers to unstressed control levels **(Fig. 2F, 2G)**. Additionally, 3369.278 markedly decreased apoptosis in stressed podocytes, as assessed by caspase 3 activity **(Fig. 2H)**.

### Hit compound increases autophagic flux and attenuates inflammation in stressed podocytes

To elucidate the MoA of 3369.278, we conducted RNA sequencing analysis of podocytes exposed to normal human sera or sera from patients with type 1 diabetes and kidney disease from the FinnDiane study, with or without 3369.278. Our findings indicate a significant reduction in the expression of TNF, pentoxifylline (PTX), chemokine (C-X-C motif) ligands (CXCLs), interleukin 6 (IL6), and interleukin 1 βeta (IL1-β), as well as in other pro-inflammatory genes known to mediate podocyte injury in DKD. Additionally, 3369.278 downregulated the expression of mitogen-activated protein kinase kinase kinase 8 (MAP3K8), a kinase critical for TNF production **(Supplementary** Fig. 1). Lipid dysmetabolism has been implicated in triggering the release of these pro-inflammatory cytokines, accelerating inflammation and contributing to kidney dysfunction^19^. In support of this, our previous work demonstrated that treatment with CD alleviated inflammation in the kidneys of mice with Alport Syndrome^11^. Building on this, our current study shows that treatment with 3369.278 reduces podocyte lipotoxicity, which likely underlies the observed reduction in inflammation, as indicated by the RNA sequencing analysis **(Table 1)**. Taken together, these findings suggest that targeting lipid accumulation and podocyte lipotoxicity may offer a promising strategy for mitigating lipid-driven inflammatory processes in DKD.

**Table 1:**
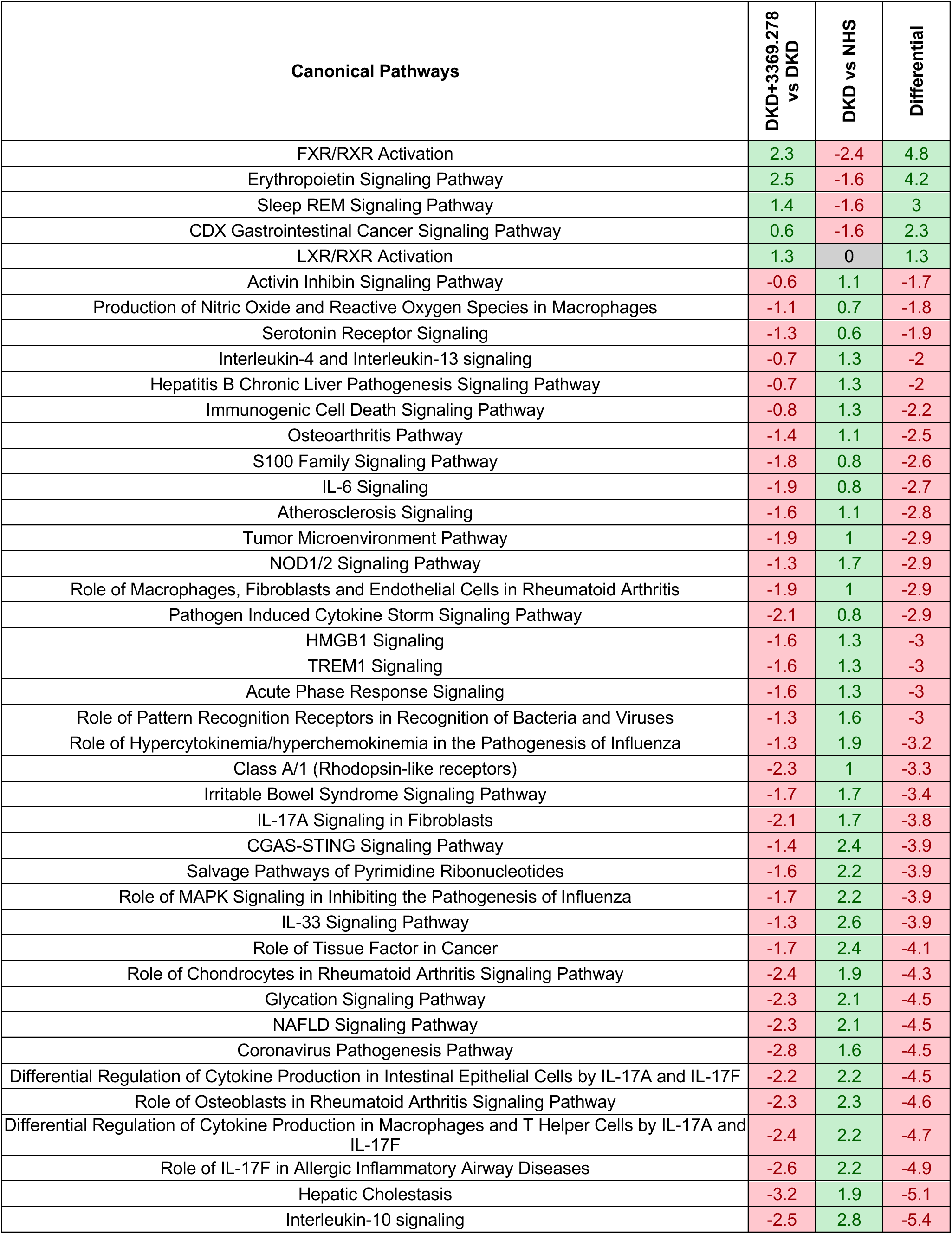
Canonical pathways modulated by 3369.278. RNA-seq data from podocytes treated with normal human sera and vehicle (NHS) or sera from patients with type 1 diabetes and kidney disease co-treated with vehicle (DKD) or 5 µM of 3369.278 (DKD + 3369.278) were analyzed using QIAGEN’s Ingenuity Pathway Analysis (IPA). The table presents the z-scores of pathways modulated by 3369.278 in comparison to DKD, and those modulated by DKD compared to NHS (p-value < 0.05, |z-score differential| > 1). Pathways upregulated by 3369.278 relative to DKD (positive differential) contribute to reduced lipid metabolism and oxidative stress. Conversely, pathways downregulated by the 3369.278 relative to DKD (negative differential) are primarily associated with inflammation.

We previously showed that promoting ABCA1-mediated cholesterol efflux attenuates proteinuria and prevents renal function decline in mouse models of proteinuric kidney disease^12^. Interestingly, 3369.278 did not restore ABCA1 expression, suggesting that its mechanism for reducing LD accumulation is independent of ABCA1 **(Supplementary** Fig. 1). Instead, we found that 3369.278 significantly upregulated the LXR signaling pathway **(Table 1)**. This is particularly noteworthy, as LXR agonists have been shown to enhance cellular cholesterol transport and reduce inflammatory responses in macrophages exposed to lipoproteins from patients with moderate to severe CKD^39^. Furthermore, in diabetic mice, LXR agonists have been reported to decrease proteinuria, renal inflammation, and renal glomerular cholesterol content by increasing ABCA1 expression^40,41^. Based on our findings, we suspect that 3369.278 may activate the LXR pathway independently of the ABCA1 pathway, thereby modulating lipid metabolism and helping to reduce inflammation.

Importantly, 3369.278 significantly reduced the expression of lysosome-associated membrane glycoprotein (LAMP), indicating potential modulation of autophagy-related mechanisms (ARMs)^42^ **(Supplementary** Fig. 1). Recent studies have highlighted the critical role of autophagy in maintaining podocyte integrity^43,44^. Dysregulated or maladaptive autophagy can lead to the accumulation of damaged cellular components and toxic substances, including in kidney cells, impairing normal cellular functions and thereby contributing to the pathogenesis of kidney diseases^45,46^. To further investigate 3369.278’s putative effect on ARMs, we assessed the levels of microtubule-associated protein light chain 3 (LC3)-positive puncta per cell, as a measure of steady-state autophagosome levels. 3369.278 significantly elevated the mean LC3+ puncta per cell **(Supplementary** Fig. 2A**, 2B)**, suggesting induced autophagy that may be due to either increased autophagosome biogenesis (suggestive of increased autophagic flux) or decreased autophagosome degradation (suggestive of reduced autophagic flux). To distinguish between these two scenarios, we assessed LC3-II levels in the presence or absence of bafilomycin A (BafA), an inhibitor of autophagosome-lysozome fusion. We found that addition of BafA following treatment with 3369.278 resulted in a strong increase in LC3-II in podocytes, demonstrating that 3369.278 induces autophagic flux in podocytes **(Supplementary** Fig. 2C, 2D, 2E).

### Preliminary hit optimization and MoA studies

We then synthesized several analogs based on the parent scaffold of 3369.278 **(Supplementary** Fig. 3) with two main objectives: 1) replacing the thiourea moiety, which could pose problems during later stages of drug development, and 2) improving aqueous solubility. These analogs were tested in our phenotypic assay, and one compound, 2726.007 **(Figure 4)** effectively attenuated LD accumulation, while restoring cell numbers to levels comparable to the unstressed control **(Fig. 5A, 5B, 5C)**.

**Figure 4:**
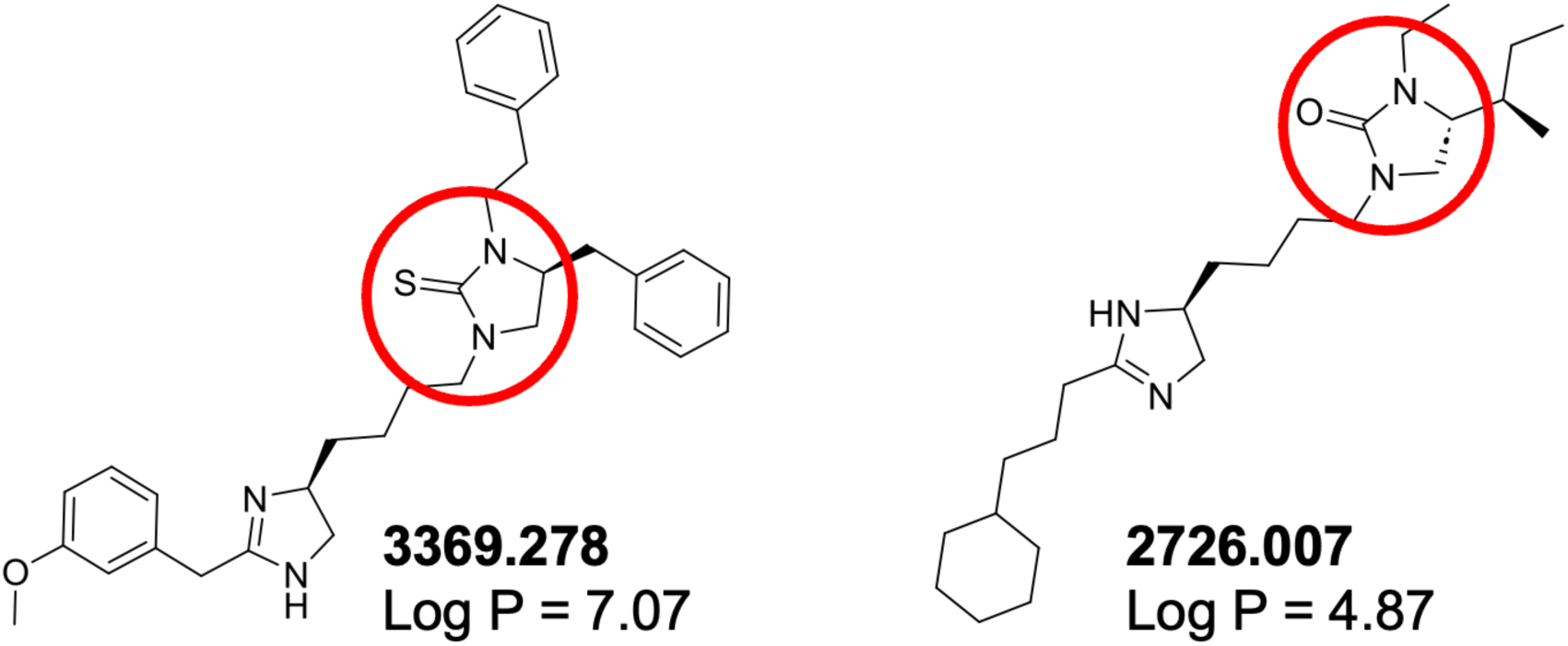
Chemical derivatization of 3369.278. 2726.007 features substantially improved aqueous solubility (lower LogP) and a urea moiety in place of the problematic thiourea.

**Figure 5:**
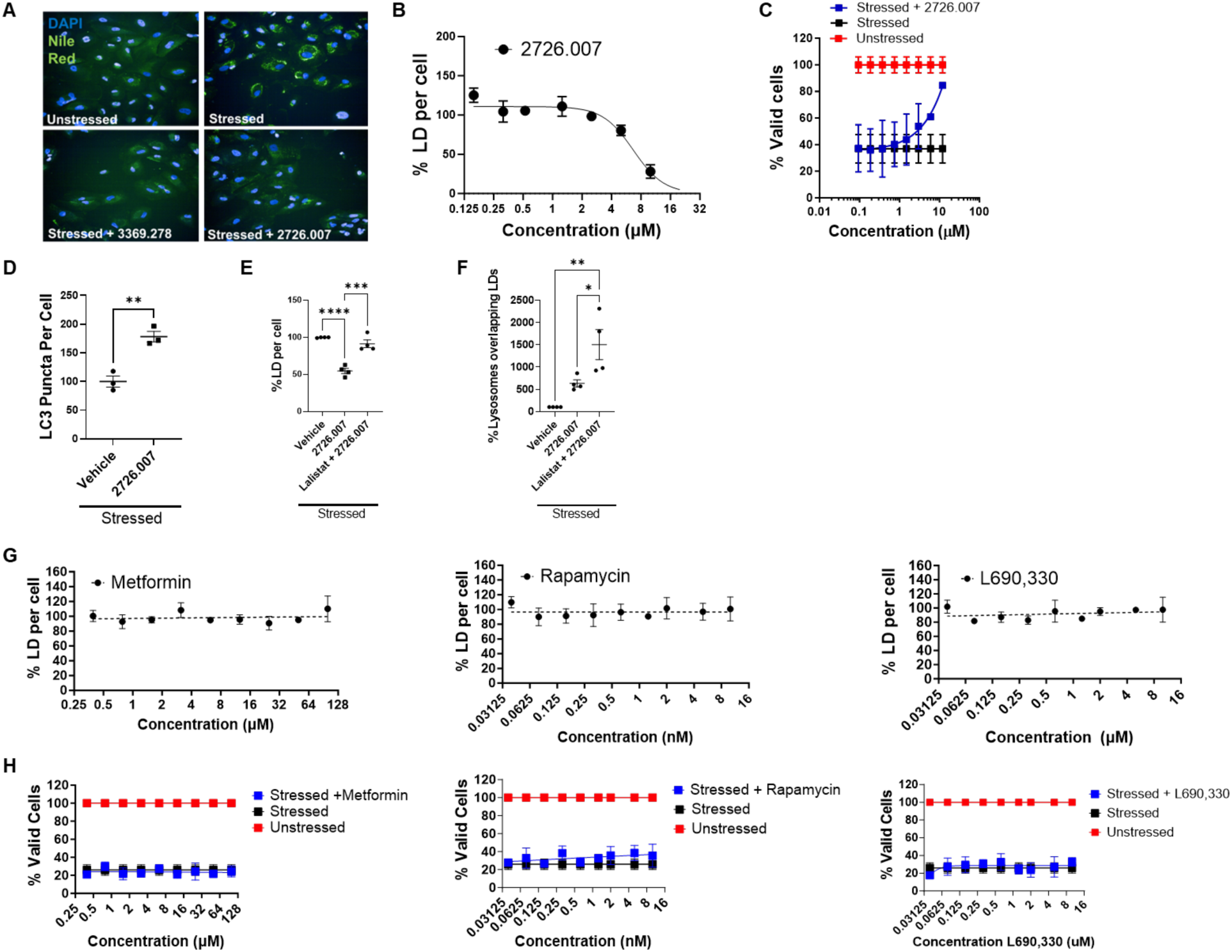
2627.007 prevents LD accumulation by inducing lipophagy. (A) Single-field OPERA confocal image of differentiated podocytes under stressed or unstressed conditions, treated with the indicated compounds or vehicle control. Cells were stained with DAPI (nuclei, blue) and Nile Red (lipid droplets, green). (B) % LDs per cell. EC50 = 6.679 µM. (C) % Valid cells. (D) LC3+ puncta numbers per cell normalized to the vehicle (100%). Data represent mean ± SEM, n=3 technical replicates. (E) % LD per cell and (F) % lysosomes overlapping LDs in stressed podocytes treated with Lalistat, an inhibitor of lysosomal acid lipase (LAL), and 2726.007 (10 µM). Data represent mean ± SEM, n=3. (G) % LD per cell and (H) % valid cells in stressed podocytes treated with metformin, rapamycin, or L690,330. Data represent mean ± SD, n=3. **p<0.01; ***p<0.001.

Given the prevalence of an imidazole moiety in adrenergic receptor (AR) agonists **(Fig. 6)**, we profiled 2726.007 in cell-based AR activity assays. Our data show that 2726.007 has no agonist activity across all three AR families and subtypes **(Table 2)**.

**Figure 6:**
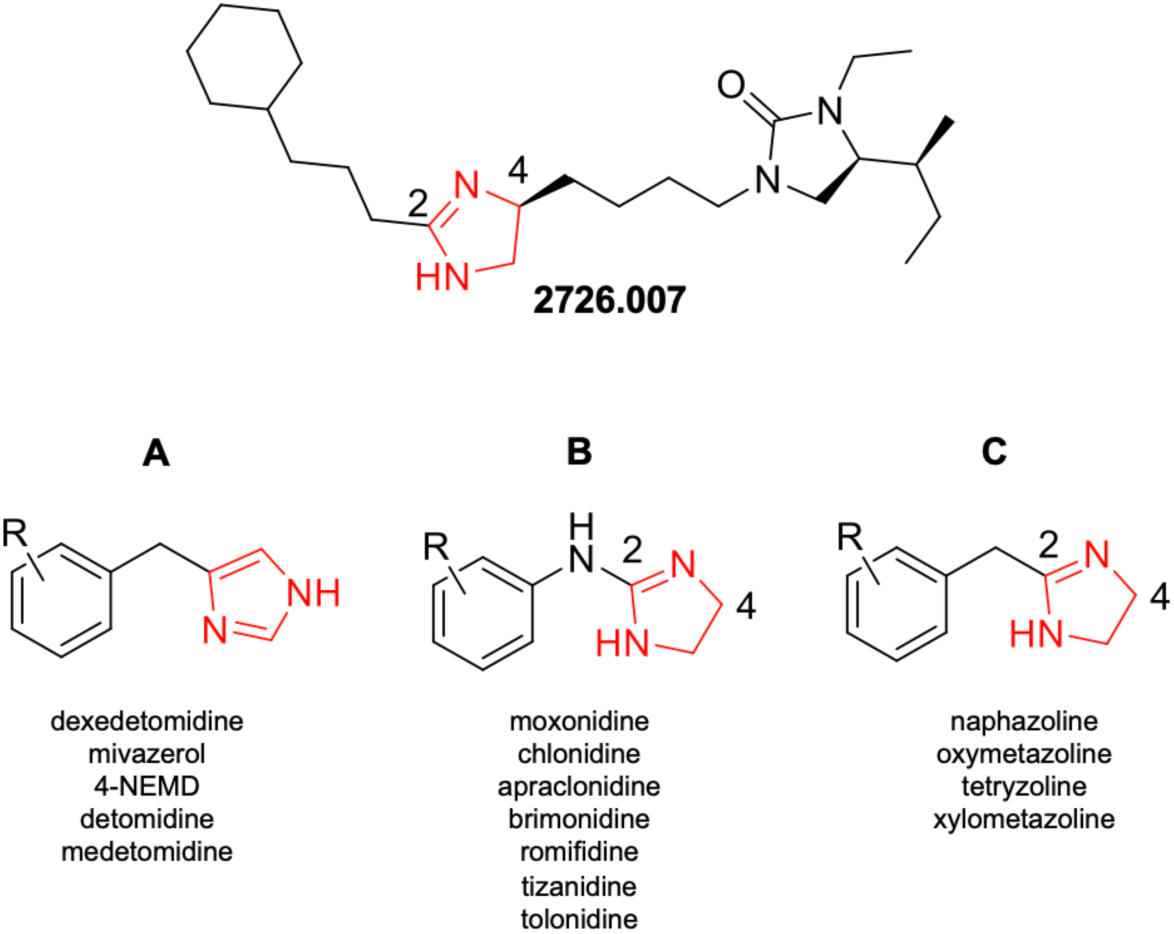
Adrenergic agonist analysis. Structures shown for 2726.007 and 3 classes of adrenergic agonists that contain an imidazole. Classes A and B are primarily alpha 2 agonists while class C is primarily alpha 1 agonists. The following features distinguish 2726.007 from known AR agonists : 1) Class A is fully aromatic and does not have a substitution at the 2 position. 2) Class B has an amine at the 2 position and no substitution at the 4 position. 3) Class C lacks a substitution at the 4 position and every adrenergic agonist of class C has an aromatic substitution at the 2 position.

**Table 2:**
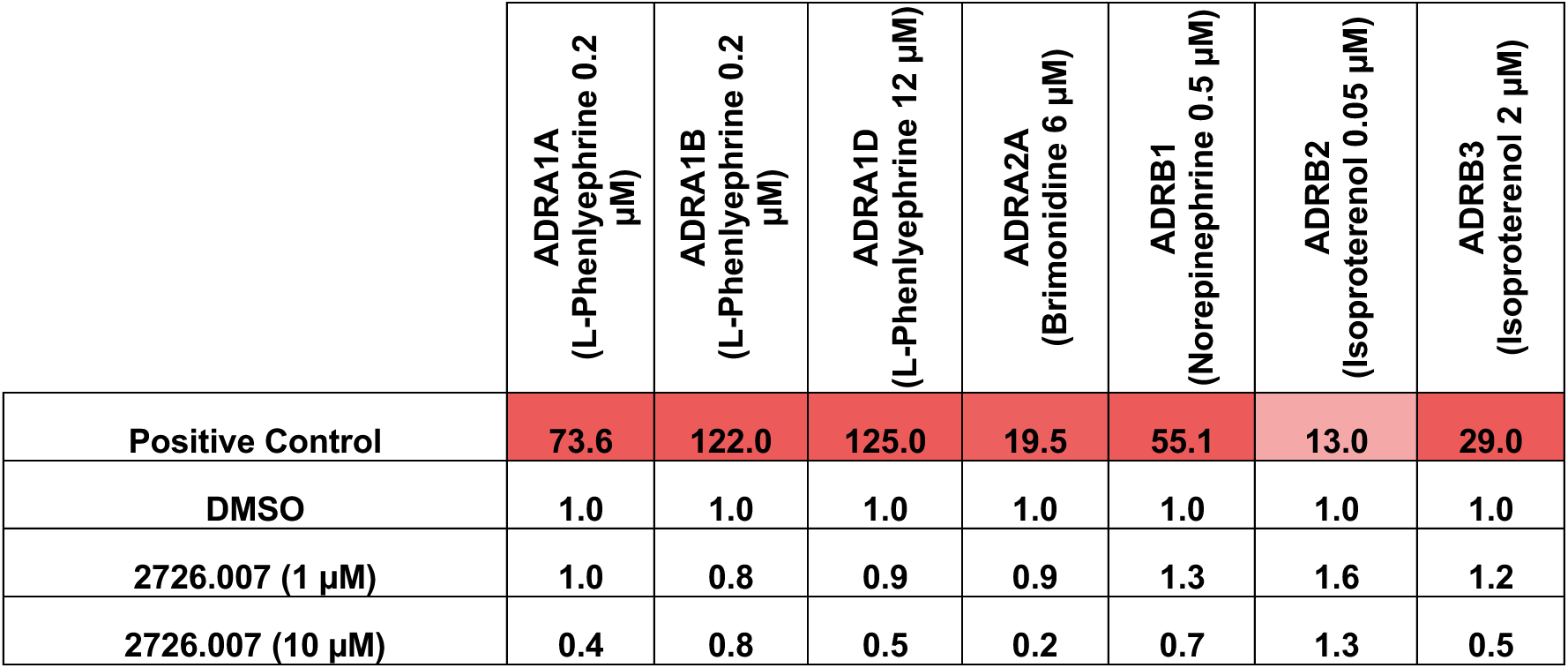
Adrenergic receptor (AR) profiling reveals no AR agonism by 2726.007. 2726.007 was tested in 7 AR reporter assays representing all three AR families. Values are mean fold-activation (n=3). The positive control for each AR and their corresponding activities are indicated for reference.

|  | ADRA1A<br>(L-Phenylephrine 0.2<br>μM) | ADRA1B<br>(L-Phenylephrine 0.2<br>μM) | ADRA1D<br>(L-Phenylephrine 12 μM) | ADRA2A<br>(Brimonidine 6 μM) | ADRB1<br>(Norepinephrine 0.5 μM) | ADRB2<br>(Isoproterenol 0.05 μM) | ADRB3<br>(Isoproterenol 2 μM) |
| --- | --- | --- | --- | --- | --- | --- | --- |
| Positive Control | 73.6 | 122.0 | 125.0 | 19.5 | 55.1 | 13.0 | 29.0 |
| DMSO | 1.0 | 1.0 | 1.0 | 1.0 | 1.0 | 1.0 | 1.0 |
| 2726.007 (1 μM) | 1.0 | 0.8 | 0.9 | 0.9 | 1.3 | 1.6 | 1.2 |
| 2726.007 (10 μM) | 0.4 | 0.8 | 0.5 | 0.2 | 0.7 | 1.3 | 0.5 |

Next, we measured LC3 puncta and found that 2726.007 significantly increased autophagosome steady-state numbers, similar to the parent 3369.278 **(Fig. 5D)**. Pathway analyses of the RNAseq data using QIAGEN’s Ingenuity Pathway Analysis (IPA) had revealed that our compounds modulate pathways regulated by Sirtuin 1 (SIRT1) and lysosomal acid lipase (LAL, gene name LIPA). Both SIRT1 and LAL play important roles in regulating lipophagy - a process in which LDs are catabolized through autophagic pathways, resulting in the lysosomal degradation of stored lipids into free fatty acids (FFAs)^1^. We hypothesized that 2726.007’s induction of ARMs leads to induction of lipophagy, thereby enabling podocytes to clear LDs and avoid lipotoxic stress. To further investigate this, we co-treated podocytes with Lalistat, an inhibitor of LAL, which blocks a critical step in the lipophagy process. Our results show that co-treatment with Lalistat blunted the effect of 2726.007 **(Fig. 5E)** and induced buildup of LD-lysosome fusions **(Fig. 5F)**, indicative of impaired clearance. These findings suggest that 2726.007 attenuates LD accumulation in stressed podocytes by inducing lipophagy. Activating lipophagy is a promising therapeutic strategy for DKD, as growing evidence highlights its crucial role in slowing disease progression^47^. A recent study showed that lipophagy was reduced in DKD, leading to increased ectopic lipid deposition (ELD) and lipotoxicity in tubular cells. Similar effects were observed in *db/db* mice and high glucose-treated HK-2 cells, where AdipoRon reduced lipid accumulation and fibrosis ^48^. To determine whether general autophagy inducers could replicate the effects of 2726.007, stressed podocytes were treated with various clinical and experimental autophagy inducers, including metformin, an AMP-activated protein kinase (AMPK) activator^49^, rapamycin, an inhibitor of mammalian target of rapamycin (mTOR)^50^, and L-690,330, a potent inhibitor of inositol monophosphatase^51^. None of these compounds reduced LD accumulation **(Fig. 5G)** or rescued podocytes from cell death in our assay **(Fig. 5H)**, suggesting that 2726.007’s MoA is distinct from the general autophagy inducers..

To investigate how 2726.007’s overall bioactivity compares to the other autophagy inducers, we used cell painting, a multiplexed high-content, high-throughput fluorescence imaging technique that extracts thousands of morphological features from cultured cells^52^. These features constitute a unique “morphological profile” for each perturbation, whereby the closer two compounds’ MoAs and molecular targets are to each other, the more similar their morphological profiles will appear. Our results show that the autophagy inducers alter the stressed phenotype in a largely similar direction within the phenotypic landscape, with rapamycin being the only one showing a substantial change. Strikingly, 2726.007 alters the phenotype in an entirely distinct direction, underscoring its unique MoA **(Fig. 7A, 7C)**. Since the traditional cell painting technique does not include a stain for lipids and is therefore highly inferential, we expanded this analysis with PhenoVue’s MultiOrganelle staining kit which includes stains for lysosomes and lipid droplets. Here, the difference between 2726.007’s MoA and the other compounds appeared much more pronounced (**Fig. 7B, 7D**), possibly due to the direct detection of changes in lipid structures.

**Figure 7:**
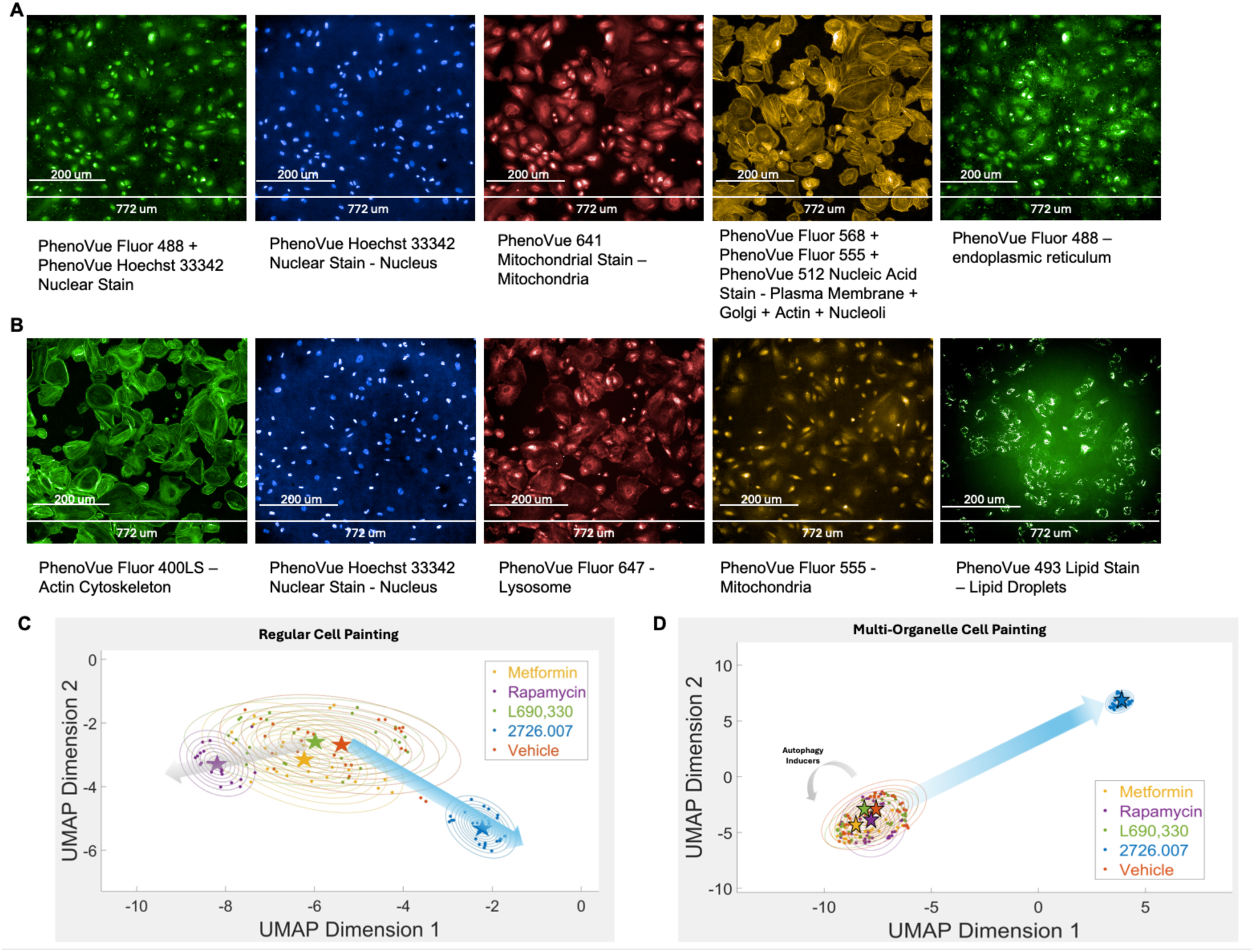
High content phenotypic profiling underscores 2726.007’s unique MoA. Cultured podocytes were stressed and treated with 2726.007 (10 µM), inducers of autophagy (Metformin (100 µM), Rapamycin (10 nM), or L690,300 (10 µM)), or vehicle following the protocol of the primary screening assay. Cultures were fixed and stained using either the (A, C) traditional Cell Painting or (B, D) Multi-Organelle kit. Plates were imaged using a Perkin Elmer Opera Phenix and >2000 phenotypic features were automatically extracted using Harmony software. UMAP was used for dimensionality reduction to conserve both local and global structure. Dots represent individual wells. The underlying Gaussian distribution is represented as contours. Asterisks denote Gaussian centers.

## CONCLUSIONS

In summary, we discovered a series of novel compounds that protect podocytes from lipotoxic injury. We demonstrate that 2726.007’s MoA involves the induction of lipophagy and appears to be mediated by the activation of the LAL, presenting it as a candidate target for DKD and other disorders of similar etiology. An advantage of this therapeutic strategy is its potential to not only halt the progression of pathological lipid accumulation but also reverse it. Given the therapeutic relevance of lipophagy to numerous human pathologies, the novel compounds reported here will serve as valuable tools for identifying molecular targets to activate cellular lipases and induce lipophagy with broad clinical impact.

## MATERIALS AND METHODS

Full details of all methods can be found in the Supporting Information.

### Cell Culture

Conditionally immortalized human podocytes (gift from Dr. Moin Saleem, University of Bristol, Bristol, England) were cultured at 33°C under permissive conditions in RPMI culture medium containing 10% fetal bovine serum (FBS), 1% penicillin/streptomycin, and 0.01 mg/ml recombinant human insulin, 0.0055 mg/ml human transferrin (substantially iron-free), and 0.005 μg/ml sodium selenite^53^. Human podocytes were thermoshifted and differentiated for 14 days at 37 °C in RPMI medium containing 10% FBS and 1% penicillin/streptomycin.

### Chemistry

The compounds utilized in this study were synthesized utilizing both a mixture-based approach and an individual compound parallel synthesis approach. The scaffold ranking and positional scanning libraires utilized in this study were all synthesized utilizing a mixture-based approach, which has been described in detail previously^27,29,30,34^. All individual compounds described in this study including 3369.273, 3369.278, and 2726.007 were synthesized using a solid phase parallel synthesis approach. Additional details are provided in the Supporting Information.

### Statistical analysis

For each statistical test, biological sample size (n) and p-value are indicated in the corresponding figure legends. The data are reported as mean ± standard error of mean or mean ± standard deviation and as individual values in the univariate scatter plots. Statistical analysis was performed using GraphPad Prism, version 9.0 (San Diego, CA, USA). Two groups were compared using independent sample t-test. Three or more groups of data were compared using one-way ANOVA followed by Tukey’s *post hoc* tests. A p-value < 0.05 was considered statistically significant.

## SUPPORTING INFORMATION

Additional experimental details, materials, and methods.

Supplementary Figure 1: Gene expression changes in podocytes stimulated with DKD patient sera and treated with 3369.278

Supplementary Figure 2: 3369.278 increases autophagic flux

Supplementary Figure 3: Preliminary hit optimization chemistry

## AUTHOR CONTRIBUTIONS

HA, MG, AF and SM supervised the project. HA, MG, and RN designed and planned the experiments. RN, BA, HD, HG, CF, CK, and JM conducted the experiments. HA and RN wrote the manuscript, and all authors edited and approved the final manuscript.

## ACKNOWLEDGMENTS

We gratefully acknowledge the Peggy and Harold Katz Family Drug Discovery Center for their continued support. We also extend our appreciation to the FinnDiane study group for their invaluable collaboration. We thank Indigo P Williams and Arianna Carrazco for her technical support. Furthermore, we acknowledge the utilization of the University of Miami Drug Discovery Core (RRID:SCR_022542). Graphical abstract was created using biorender.

