## Supplementary Data for "Novel Lipophagy Inducers as Potential Therapeutics for Lipid Metabolism Disorders"

**SUPPLEMENTARY MATERIAL TABLE OF CONTENTS**

Supplementary Materials and Methods

Supplementary Figure 1: Gene expression changes in podocytes stimulated with DKD patient sera and treated with 3369.278

Supplementary Figure 2: 3369.278 increases autophagic flux

Supplementary Figure 3: Preliminary hit optimization chemistry

### Supplementary Materials and Methods

### Compound Purification and Characterization

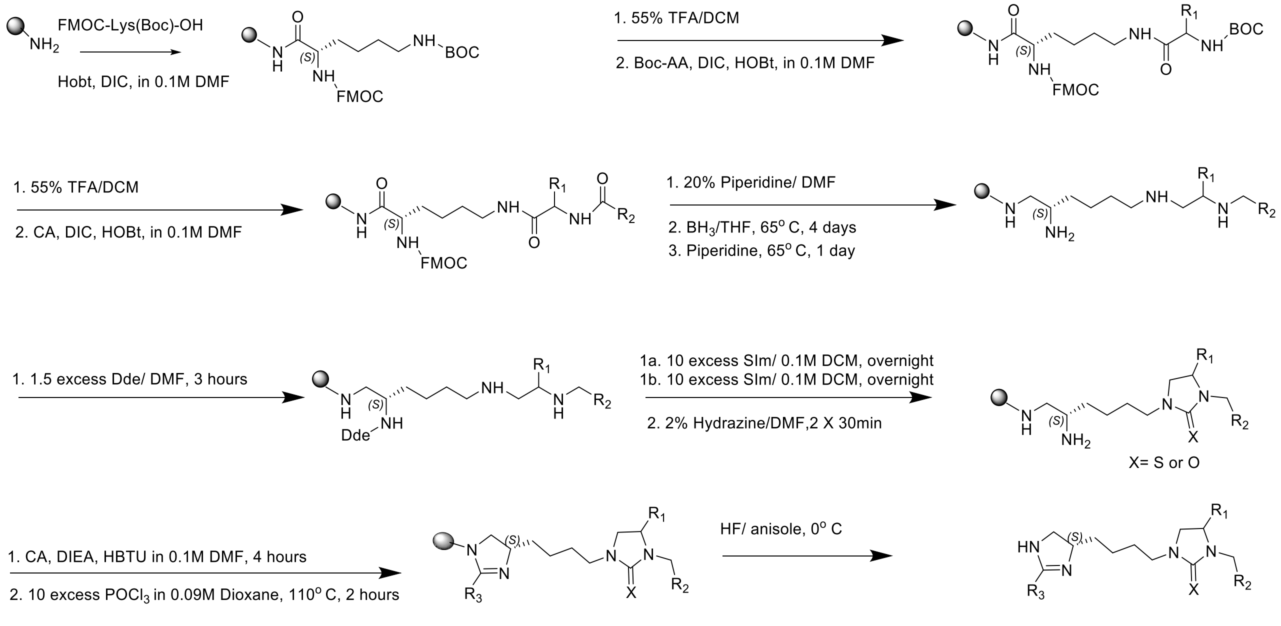


*Scheme 1. Synthetic scheme for compounds of the 3369 and 2726 series.*

All reagents were commercially available and used without further purification. The compounds were synthesized as described in **Scheme 1** above using standard Boc chemistry and previously reported[34]. The solid phase synthesis was performed using the "tea-bag" methodology[54]. The final compounds were purified using preparative HPLC with a dual pump Shimadzu LC-20AB system equipped with a Luna C18 preparative column (21.5 x 150 mm, 5 micron) at λ = 214 nm, with a mobile phase of (A) $H_2O$ (+0.1% formic acid)/(B) acetonitrile (ACN) (+0.1% formic acid) at a flow rate of 15 mL/min; gradients varied by compound based on hydrophobicity. Molecular Weight and Log P values were calculated utilizing ChemDraw Version 22.2.0.

**3369.273 (4S)-4-benzyl-3-phenethyl-1-(4-((5S)-2-(2-phenylpropyl)-4,5-dihydro-1H-imidazol-5-yl)butyl)imidazolidine-2-thione.** 3369.273 was synthesized using Scheme 1 and the following reagents: (100mg) MBHA resin starting material, Boc-L-Phe-OH ($R_1$), Phenylacetic acid ($R_2$), 3-Phenylbutyric acid ($R_3$), and X = S. The final crude product was purified using HPLC as described above, $C_{34}H_{42}N_4S$, Molecular Weight 538.8, Log P 7.94.

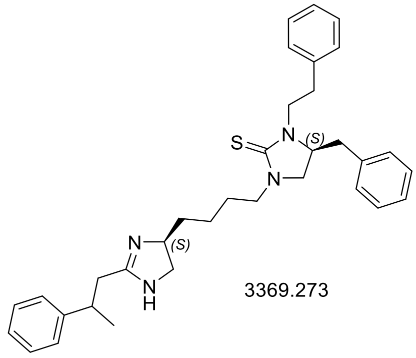


3369.273

**3369.278 (S)-4-benzyl-1-(4-((S)-2-(4-methoxybenzyl)-4,5-dihydro-1H-imidazol-5-yl)butyl)-3-phenethylimidazolidine-2-thione.** 3369.278 was synthesized using Scheme 1 and the following reagents: (100mg) MBHA resin starting material, Boc-L-Phe-OH ($R_1$), Phenylacetic acid ($R_2$), 3-Methoxyphenylacetic acid ($R_3$), and X = S. The final crude product was purified using HPLC as described above. $C_{33}H_{40}N_4OS$, Molecular Weight 540.8, and Log P 7.07.

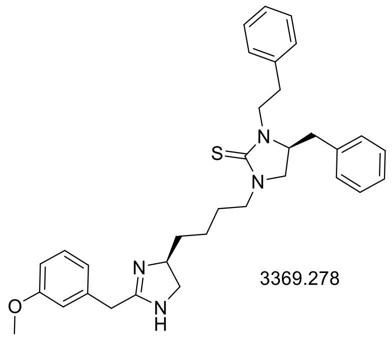


3369.278

**2726.007 (R)-4-((R)-sec-butyl)-1-(4-((S)-2-(3-cyclohexylpropyl)-4,5-dihydro-1H-imidazol-5-yl)butyl)-3-ethylimidazolidin-2-one.** 2726.007 was synthesized using Scheme 1 and the following reagents: (100mg) MBHA resin starting material, Boc-D-Ile-OH ($R_1$), Acetic Acid ($R_2$), Cyclohexanebutyric Acid ($R_3$), and X = O. $C_{25}H_{46}N_4O$, Molecular Weight 418.7, Log P 4.87.

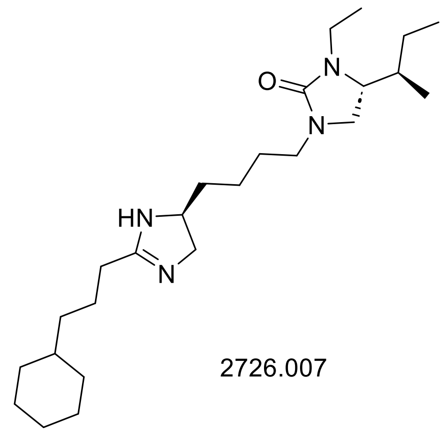

2726.007

**Counter-screening in adrenergic receptor assays**

Counter-screening for adrenergic receptor (AR) activity was performed using INDIGO INC's cell-based AR reporter assays (#IB31002, #IB31102, #IB31202, #IB34002, #IB32002, #IB32102, #IB32202), which employ luciferase-based functional assays to measure AR activation. 2726.007 was tested at concentrations of 1 μM and 10 μM across seven AR reporter assays, representing all three AR families: ADRA1A, ADRA1B, ADRA1D, ADRA2A, ADRB1, ADRB2, and ADRB3. A positive control was included for each AR subtype: L-phenylephrine (0.2 μM) for ADRA1A and ADRA1B, L-phenylephrine (12 μM) for ADRA1D, brimonidine (6 μM) for ADRA2A, norepinephrine (0.5 μM) for ADRB1, isoproterenol (0.05 μM) for ADRB2, and isoproterenol (2 μM) for ADRB3.

**Lipid Droplet Assay**

Conditionally immortalized human podocytes were seeded in 96-well plates at a density of 6,000 cells per well. At day 12 of differentiation, the podocytes were treated with the FIU-CTS mixture library at four different concentrations (5, 1, 0.2, and 0.04 μg/mL) or 2-fold serial dilutions of 3369.273, 3369.278, 2726.007, metformin (#317240 - Millipore Sigma), rapamycin (553210 - Millipore Sigma)), or L690,330 (HY-101075 - MedChemExpress) and incubated for 1 hour at 37°C in 5% CO2. After the 1-hour incubation, puromycin (30 μg/mL) (Caymen Chemical, 13884) or sera from patients with type 1 diabetes from the Finnish Diabetic Nephropathy study were added as inducers of podocyte stress, and the cells were incubated for an additional 48 hours at 37°C in 5% CO2. Following the incubation, the cells were fixed at room temperature for 20 minutes by adding paraformaldehyde (4%) and sucrose solution (4%) in phosphate-buffered saline (PBS). After fixation, the cells were stained with DAPI (1 mg/mL at final concentration; 1:5000) (ThermoFisher Scientific, #62248), Nile Red (200 μM in 100% methanol at final concentration; 1:1000) (Invitrogen, #N1142), Alexa Fluor 647 Phalloidin (1200 U/mL in DMSO at final concentration; 1:1000) (Invitrogen, #A22287), and HCS CellMask Deep Red 650/655 (10 mg/mL in DMSO at final concentration; 1:80,000) (Invitrogen, #H32712) in PBS at room temperature for 30 minutes. The plates were then washed with PBS and imaged on an Opera Phenix high-content screening microscope (20X water objective, 36 fields per well). Images were acquired with three channels: UV 365 for the nuclear stain (DAPI), 488 channel for the Nile Red LD stain, and 647 channel for the Phalloidin and CellMask stain. The images were analyzed using Perkin-Elmer's Harmony software and outcome measures for LDs, LD intensity, valid cell counts, and z-factor were calculated for each well as previously described[18]. P[

**Caspase-3 Activity Assay**

Caspase-3 activity was assessed in differentiated human podocytes under stressed or unstressed conditions, treated with 3369.278 (starting at 10 μM with serial 1:2 dilutions) or vehicle control for 48 hours. The assay was performed using the caspase-3 assay kit (Abcam, cat#ab39401) according to the manufacturer's instructions.

**LC3 Puncta**

Conditionally immortalized human podocytes were seeded in 96-well plates at a density of 6,000 cells per well. At day 12 of differentiation, podocytes were treated with 2726.007 (10 μM) or 3369.279 (10 μM) and incubated for 1 hour at 37°C in 5% CO2. After the 1-hour incubation, puromycin (30 μg/mL) was added as inducer of podocyte stress, and the cells were incubated for an additional 48 hours at 37°C in 5% CO2. Following the 48-hour incubation, cells were fixed at room temperature for 20 minutes by adding paraformaldehyde (4%) and sucrose solution (4%) in phosphate-buffered saline (PBS), and then permeabilized for 10 minutes at room temperature with 0.3% Triton X-100 in PBS. A blocking step was performed using 5% BSA and 0.3% Triton X-100 in PBS for 1 hour at room temperature, followed by incubation with Rabbit Polyclonal anti-LC3B Antibody (Novus Biologicals NB600-1384, 1:1000) at 4°C overnight. Alexa Fluor 488-conjugated secondary antibody was used for 1 hour at room temperature. Podocytes were then stained for segmentation using DAPI (1 mg/mL at final concentration, 1:5000) (ThermoFisher Scientific, #62248), Alexa Fluor 647 Phalloidin (1200 U/mL in DMSO at final concentration, 1:1000) (Invitrogen, #A22287), and HCS CellMask Deep Red 650/655 (10 mg/mL in DMSO at final concentration, 1:80,000) (Invitrogen, #H32712) in PBS at room temperature for 30 minutes, followed by imaging on an Opera Phenix high-content screening microscope (20X water objective, 36 fields per well). The images were analyzed using PerkinElmer's Harmony software.

**Lipid Droplet Lysosome Assay**

Conditionally immortalized human podocytes were seeded in 96-well plates at a density of 6,000 cells per well. At day 12 of differentiation, podocytes were treated with 2726.007 (10 μM) in the presence of absence of Lalistat (1 μM) (SML2053, Millipore Sigma) and incubated for 1 hour at 37°C in 5% CO2. After the 1-hour incubation, puromycin (30 μg/mL) was added as inducer of podocyte stress, and the cells were incubated for an additional 48 hours at 37°C in 5% CO2. Following the incubation, lysotracker Deep Red (75 nM at final concentration; Invitrogen, cat #: 2426290) was added to podocytes and the plate was incubated for 1 hour at 37°C in 5% CO2. After 1 hour-incubation, cells were fixed at room temperature for 20 minutes by adding paraformaldehyde (4%) and sucrose solution (4%) in phosphate-buffered saline (PBS), and then stained with DAPI (1 mg/mL at final concentration; 1:5000) (ThermoFisher Scientific, #62248) and Nile Red (200 μM, in 100% methanol at final concentration; 1:1000) (Invitrogen, #N1142) in PBS at room temperature for 30 minutes. The plates were then washed with PBS and imaged on an Opera Phenix high-content screening microscope (20X water objective, 36 fields per well). Images were acquired with three channels: UV 365 for the nuclear stain (DAPI), 488 channel for the Nile Red LD stain, and 647 channel for the lysotracker dye. The images were then analyzed using Perkin-Elmer's Harmony software and outcome measures for LDs, lysosomes overlapping LDs were calculated for each well as previously described[18].

**Cell Painting**

Conditionally immortalized human podocytes were seeded in 96-well plates at a density of 6,000 cells per well. At day 12 of differentiation, the podocytes were stressed and treated with 2726.007 (10 μM), metformin (100 μM) (#317240 - Millipore Sigma), rapamycin (10 nM) (553210 - Millipore Sigma)), L690,330 (10 μM) (HY-101075 - MedChemExpress) or vehicle following the protocol of the primary screening assay. The cells were then fixed and stained using either the PhenoVue Cell Painting Kit (#PING11 - Revvity) or the PhenoVue Multi-Organelle Staining Kit (#PMOS11 - Revvity) according to the manufacturer's instructions. Plates were imaged using a

Perkin Elmer Opera Phenix and more than 2000 phenotypic features were automatically extracted using Harmony software. Dimensionality reduction in MATLAB was performed to visualize the findings in 2D pseudospace, where the proximity between treatments reflects the similarity of their MoAs.

**RNA Sequencing Analysis**

Conditionally immortalized human podocytes were cultured and differentiated as previously described[53]. On day 12 after thermoshifting, differentiated podocytes were stimulated with 4% v/v normal human sera (NHS) co-treated with the vehicle, or DKD patient sera co-treated with either vehicle or 3369.278 (5 μM) for 48 hours (3 replicates per treatment per serum condition, 12 total). After 48 hours of treatment, RNA was extracted and processed for RNA sequencing using the PureLink RNA Mini Kit (Invitrogen, 12183018A). Total RNA was prepped with the Tecan Universal Plus mRNA-Seq with NuQuant (M01485 v6) using 400ng via Qubit and 14 cycles PCR. Libraries were sequenced on the Illumina Novaseq X Plus and >48M PE150 were generated per sample. Reads from RNA-Seq were mapped to reference genome GRCh38 using STAR (ver.2.5.0) aligner[55]. Raw counts were generated based on Ensembl gene models (GENCODE ver.21) with featureCounts (ver.1.5.0)[56]. Differential expression genes (DEGs) were identified using DESeq2[57], with significance corrected for multiple hypothesis testing (FDR<0.05). Pathway analyses of the RNAseq data was performed using QIAGEN's Ingenuity Pathway Analysis (IPA) (Ingenuity Systems, Redwood City, CA, USA).

**Protein Extraction and Western Blot Analysis**

To assess autophagic flux, podocytes were treated with 3369.278 for 18 hours, followed by Bafilomycin (10nM) (Cayman Chemical; Cat # 11038) for 6 hours. Podocytes were then homogenized in ice-cold RIPA Lysis and Extraction Buffer (ThermoFisher Scientific) with protease inhibitor (Pierce™ Protease Inhibitor Tablets, EDTA-free ThermoFisher Scientific A32965) and phosphatase inhibitor (PhosStop EASYpack Sigma Ref: 04 906 845 001). Protein concentration was quantified using the Pierce BCA Protein Assay kit according to the manufacturer's protocol. Samples were prepared in 4× Laemmli buffer, and 20 µg of protein was loaded onto 4%–20% SDS-polyacrylamide gel electrophoresis (SDS-PAGE) gels (Bio-Rad, Hercules, CA) and transferred to Immobilon-P PVDF membranes (Bio-Rad, Hercules, CA). Membranes were blocked in 5% BSA skim milk for 1 hour at room temperature followed by incubation with rabbit polyclonal anti-LC3B antibody (Novus Biologicals NB600-1384 1:1000) at 4°C, overnight. After washing, membranes were incubated with anti-rabbit IgG-HRP antibodies (Promega, 1:10,000). Signal was detected with Radiance ECL (Azure, Dublin, CA) using Azure c600 Imaging System.

### Supplementary Figures

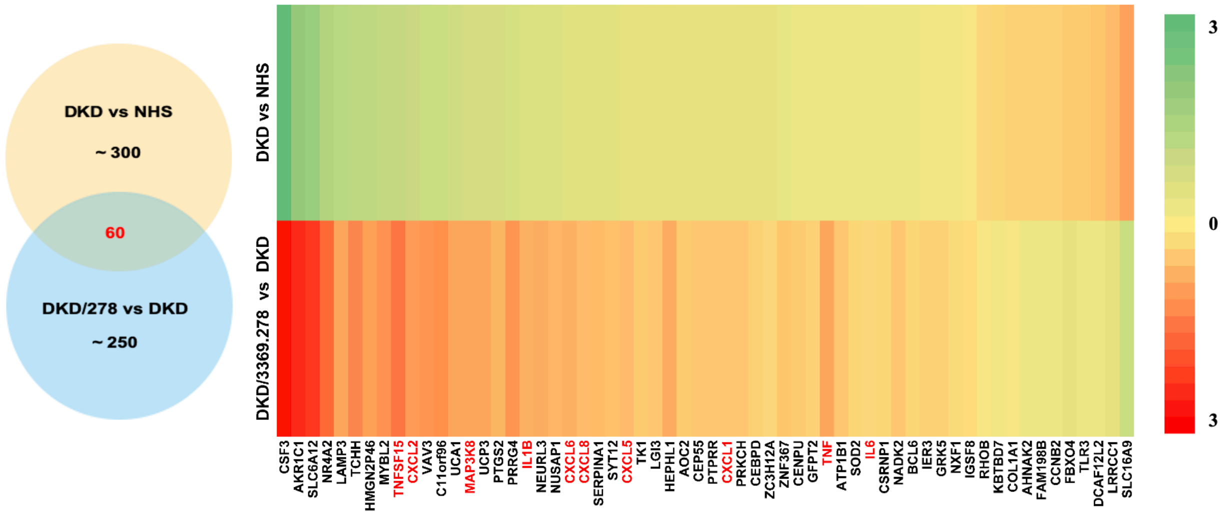


**Supplementary Figure 1: Gene expression changes in podocytes stimulated with DKD patient sera and treated with 3369.278.** Podocytes were stimulated with 4% v/v normal human sera (NHS) co-treated with the vehicle, or DKD patient sera co-treated with either vehicle or 3369.278 (5 µM), for 48 hours. RNAseq analysis showed that 60 differentially expressed genes were shared between the "DKD vs. NHS" and the "DKD/3369.278 vs. DKD" comparative analyses (data represent log2 of mean fold-change). Remarkably, 3369.278 reversed the direction of expression change induced by exposure to DKD patient sera for each of those genes.

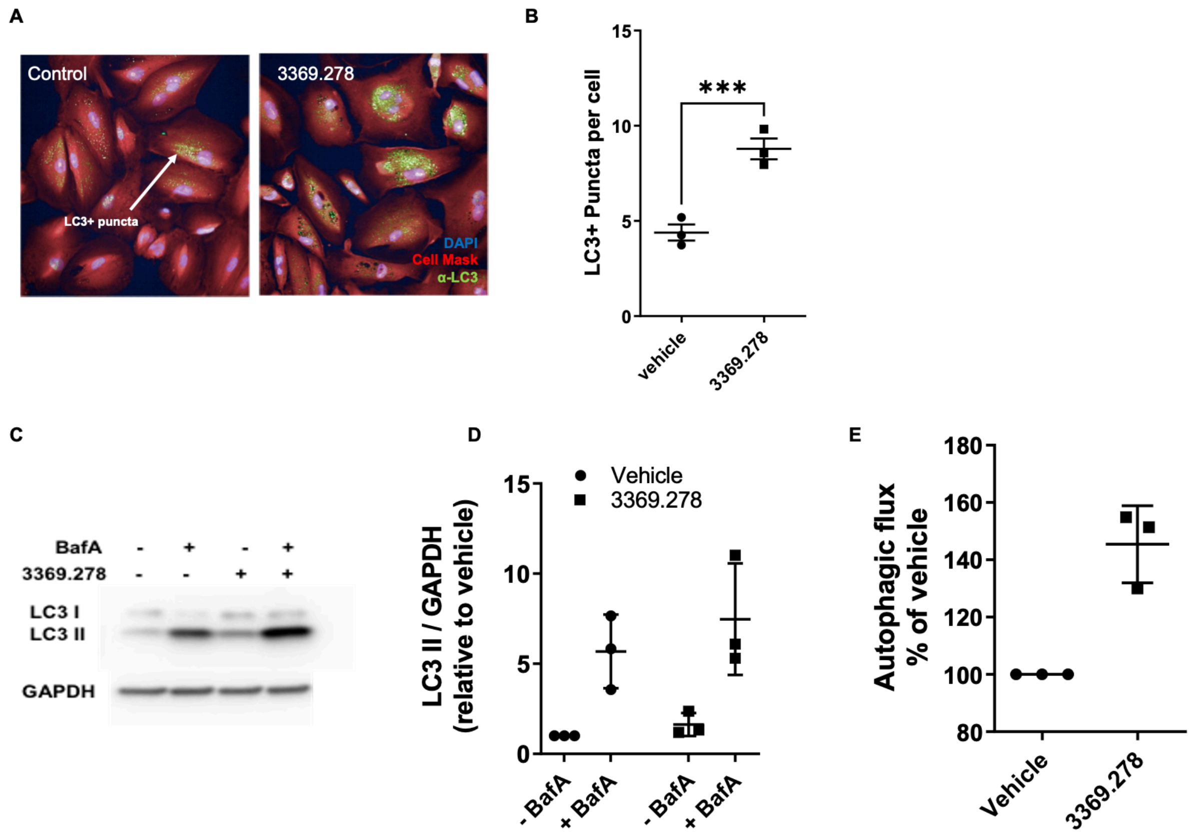


**Supplementary Figure 2: 3369.278 increases autophagic flux.** (A) Single-field OPERA confocal image of differentiated podocytes under stressed conditions treated with 3369.278 (10 µM) for 18 hours or vehicle control. Cells were stained with DAPI (nuclei, blue), CellMask Deep Red (cell bodies, red), and α-LC3 (green). (B) LC3+ puncta numbers per cell normalized to the vehicle (100%). Data represent mean ± SEM, n=3. ***p<0.001. (C) Analysis of Western blot for LC3II and GAPDH in differentiated podocytes under stressed conditions treated with 3369.278 (10 µM) for 18 hours or vehicle control, followed by Bafilomycin (10nM) for 6 hours or without bafilomycin. (D) Densitometric analysis of LC3II/GAPDH. Data represent mean ± SD, n=3 technical replicates. (E) Autophagic flux, calculated as the difference between +BafA and -BafA levels in LC3II, and normalized to the vehicle (100%).

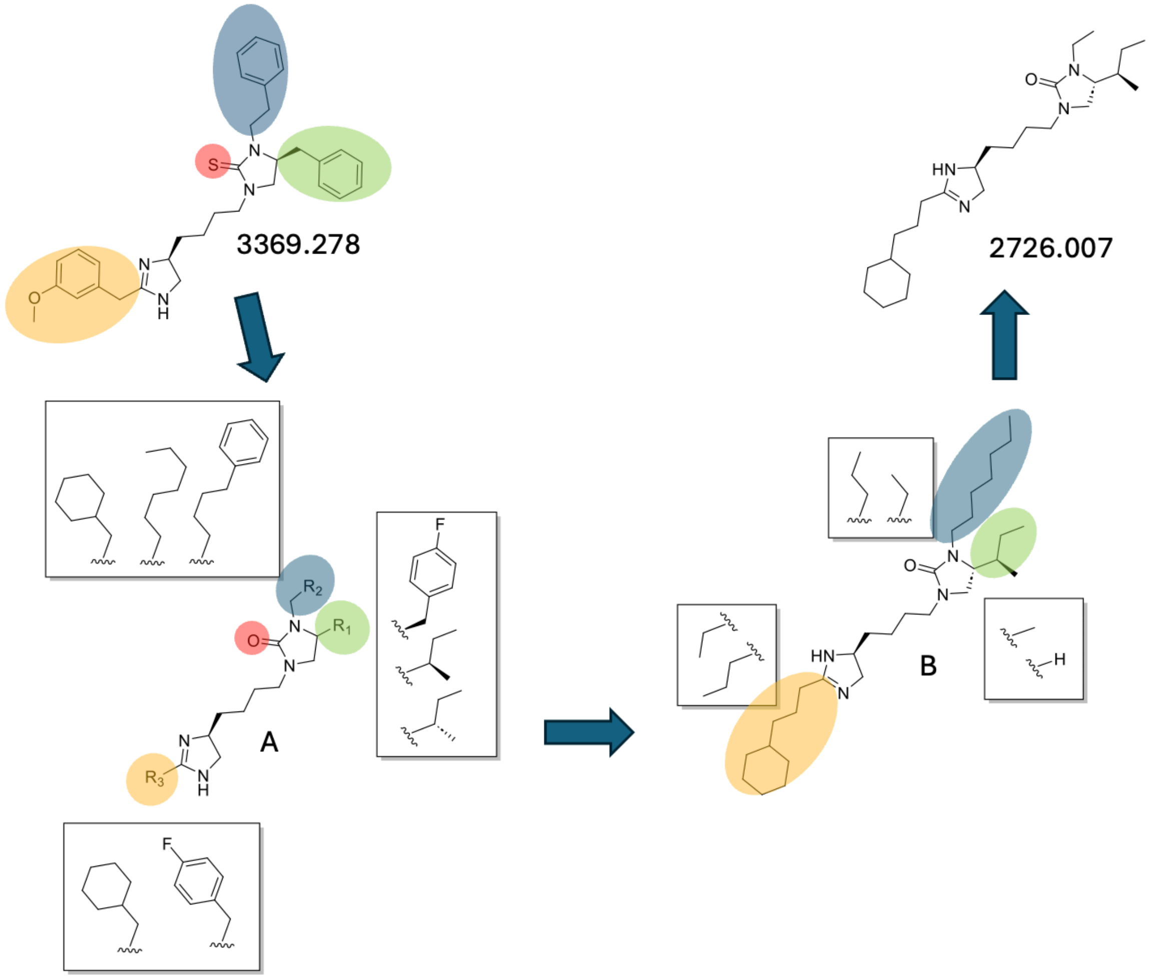


**Supplementary Figure 3: Preliminary hit optimization chemistry.** We synthesized several analogs based on the parent scaffold of 3369.278 with two main objectives: replacing the thiourea moiety, which could pose problems during later stages of drug development, and improving aqueous solubility to enable in vivo use. Starting with our initial best hit 3379.278 we made a set of 18 compounds around the urea scaffold A. The figure shows the 3 functionalities utilized at $R_1$, the three at $R_2$, and the two functionalities at $R_3$ (boxed functionalities next to blue, orange, and green on A). The 18 compounds come from making all combinations of the R functionalities (3x3x2=12). The functionalities were chosen based on previous phenotypic screening data of the 3369 positional scanning library (data not shown). After screening the 18 compounds in the phenotypic assay we found a new hit compound B where we had effectively removed the thiourea moiety and now had a scaffold with a urea. To increase the solubility of the hit compound B we made a series of single substitution analogs where we systematically truncated the three diversity positions (boxed functionalities next to blue, orange, and green on B). These single substitution analogs all had significantly lower LogP values. After screening these compounds in the phenotypic assay 2726.007 was identified as an improved hit compound.
